# Novel determination on root-knot nematodes; microspace, mineral profile and transduction of metabolic signals in tomato

**DOI:** 10.1101/851832

**Authors:** Víctor García-Gaytán, Esteban Sánchez-Rodríguez, José J. Ordaz-Ortiz, Josaphat M. Montero-Vargas, Olimpia Alonso-Pérez, Luis Rojas-Abarca, Elena Gómez-Cabrera, Juan L. Negrete-Guerrero

## Abstract

Globally, nematodes are parasites that destroy many crops, however, their presence also serves as an indicator of the state of soil health. Tomato is a very studied, cultivated vegetable. In the micro-space of the knots we determine its pathogenesis. The macronutrients, micronutrients and beneficial elements present on root-knot nematodes were C>O> N (54.73, 37.57, and 3.55%), phosphorus (0.133%), Na>Ca>Mg>K cations (1.13, 1,032, 0.094, and 0.048%), S (0.247%), micronutrients Fe>Cl (0.163, 0.027%), and beneficial elements Si>Al (0.354, 0.985%). Progenesis of the chromatograms (ES+/ES-), 202 compounds were detected in the first polarity, in which 195 have at least one possible identification candidate, the second were 42 compounds, and 41 match with at least one compound. Putative compounds with the highest scores are reported in this research: kukoamina (57.6), furmecyclox (51.1), feruloylputrescina (55.3), N1-trans-feruloylagmatine (53.9), dehydrotomatin (49.8, 50.8, 52.3, 52.6), jurubine (54.8,51.6), etnangien (50.2), dehydromelilotoside (41.5), tomatine (55), minutissamide (51.7). Nematodes in state (J2), six nematodes within the knots with an average length of 1.16 mm, were found and their interaction with other microorganisms is possible. With the high concentrations of Na^+^ in the root, the concentration of the cation Mg^2+^ and K^+^ decreases; while the N does not present any change.

## Introduction

For plants to express their maximum yield potential, proper management of root nutrition, stimulation and bioprotection is necessary (García-Gaytán et al., 2018a), the plant genotype and an adequate nutrient solution (García-Gaytán et al., 2018b), foliar nutrient applications (García-Gaytán et al., 2013), and environmental monitoring (prevention of pests and diseases) (García-Gaytán et al., 2016). Although, tomato cultivation is one of the most studied and cultivated in different growth conditions, it is also a vegetable of greater demand in national and international markets. These innovations and value added must be put in place for other crops, such as native species of great gastronomic recognition (García-Gaytán *et al*., 2017).

The improvement in tomato yield is also due, thanks to hybrids developed in the mid-1980s (Grandillo *et al*., 1999). An integrated control management makes a significant contribution to agricultural sustainability and environmental quality. Organic matter (OM) and practices that increase total microbial activity in the soil improves pathogen suppression by increasing competition for nutrients (Ghorbani *et al*., 2009). Compost reduces disease development, due to the activity of consortia of antagonistic microorganisms (Hadar and Papadopoulou, 2012). In addition, crop rotation increases bacterial populations, and soil microbial activity (Larking *et al*., 2010). Root-knot nematodes (RKN) form a feeding site by inducing several individual cells to become giant cells (Hoth *et al*., 2008). These parasitic nematodes destroy 12.3% of crop production annually, which translates to more than 157 billion dollars worldwide (Hassan *et al*., 2013). While nematodes are active in the soil throughout the year, it therefore has the potential to provide a holistic measurement of the biotic and functional state of soils (Ritz and Trudgill, 1999). Studies of wave movements in Newtonian fluids (viscosity) of the *Caenorhabditis elegans* nematode have been carried out, the wave march continuously varies with changes in the external load, that is, as the load increases, the wavelength and frequency decrease wave (Fang-Yen *et al*., 2010). Previously, it had been suggested that metabolic engineering in plants would allow a critical evaluation of the role of individual compounds in plant-rhizosphere communication (Van Dam and Bouwmeester, 2016). The term microcosm has also been discussed to study the diversity of plants and their effect on the biomass of fungi and soil bacteria (Eisenhauer *et al*., 2017), and explain the beneficial effects of the interaction of species of plants, and worms on nematodes (Niu *et al*., 2019). This research, the term micro-space is addressed for the first time. We demonstrate the number of nematodes that can accommodate a knots (pathogenesis) and corroborates the wave movement in it. We also performed the analysis of macronutrients, micronutrients and beneficial elements on (RKN), metabolic compounds on (RKN). And the putative compounds with the highest score. To our knowledge, this is the first article worldwide that addresses the wave movement of the nematode in the microspace of the knots - elementary content - metabolomic profile (compounds of great agronomic relevance and for future research).

## Material and Methods

### Biological material and micro-environment conditions

Roots infested by nematodes (*Meloydogine* ssp.) in tomato plants (*Solanum* ssp.) were collected in a greenhouse in Guanajuato, Mexico. The conditions inside the greenhouse were; CO_2_ concentration was 280 ppm, air temperature 88 °F and relative humidity (RH) of 49% (Temp/RH/CO_2_, Hand-Held Meter, Spectrum^®^ Technologies, Inc.).

### Scanning electron microscopy / energy dispersive X-ray spectrometry (SEM-EDS) Scan pathogenesis: Micro-space over the knots

Root-knot nematodes, the knots were deposited in 50 mL-1 falcon-type plastic tubes with distilled water. Root knots was removed and immersed in liquid nitrogen at −195.79 °C. A cross section was made to the tissue. With the use of a scanning electron microscopy/X-ray spectrometry of energy dispersion (SEM-EDS) micrographs of the nematodes were taken into the gill and their dimensions were determined (μm). The nematodes were also observed in a stereoscopic microscope (Leica, EZ4HD, 8-35x) Supplementary Fig. S1.

### Elementary profile on (RKN)

To identify the elementary composition and the relative distribution of the primary, secondary, micronutrient, and beneficial elements on RKN. The samples were processed according to the methodology of García-Gaytán et al. (2019). The determination of the elementary content was determined directly on the knots. The relative content was determined in a scanning electron microscope (Scanning Electron Microscope, Model 7582, England), the value of the elementary composition corresponds to the average of five repetitions in the tissue. They were plotted with GraphPad Prism 7.0 software (GraphPad Software, Inc., La Jolla, CA, USA).

### LC-MS Q-TOF metabolomic profile on (RKN)

The knots were deposited in two 50 mL^−1^ falcon plastic tubes in the first tube containing FAA fixative solution (formaldehyde, alcohol, distilled water) in a second tube the gills contained HPLC water. The samples of each tube were processed with liquid nitrogen - 195.79 °C. The fine powders of the samples were deposited in eppendorf tubes each. The samples were subsequently centrifuged, to separate the supernatant from the tissue. With a miVac the extracts were dried and reconstituted with 250 μL equal to the initial gradient of liquid chromatography. The extracts were run by LC-MS Q-TOF in both polarities for a m/z range of 50-1500.

### Liquid chromatography

An ACQUITY UPLC® HSS T3 1.8 μm 2.1 × 100 mm column was used for liquid chromatography. In mobile phase A, H_2_O:0.1% FA was used and for mobile phase B, 0.1% FA ACN was used, with a flow rate of 0.4 mL/min. With six time gradients (min): initial, 0.50, 25, 26.50, 26.75 and 30 respectively. The injection volume was 10 μL, with a column temperature of 40 ° C.

### Mass spectrometry (MS)

For the MS, a Water Synapt G1 Q-TOF device was used, with polarity (ES+/ES-), capillary (kV) 3.0 and 2.6, sampling cone (V) 40.0, extraction cone (V) 4.0, source temperature (°C) 120, desolvation temperature (°C) 350, desolvation gas flow (L/h) 500, Scan time (s) 1.5, LM resolution 4.7, HM resolution 15, trap CE 6.0, transfer CE 4.0, source (mL/min) 0.0, trap (mL/min) 1.5, detector 1850 and 1750, multiplier 650, finally the autosampler temperature was 6 ° C.

## Results

### Pathogenesis

Root-knot nematodes were identified male nematodes in juvenile state within the knot. In fig. 2, it can be seen that the nematodes show wave mobility in the micro-spaces (communication galleries) within the knot (Fig. 2A). Cross section of the nematode within the knot micro-space, with wave mobility (Fig. 2B). We demonstrate, for the first time, that, in an infested gall, more than six nematodes can be housed (Fig. 2C). The nematodes have an elongated cylindrical shape with a length of 1.16 mm (Fig. 2D).

**Fig. 2.**
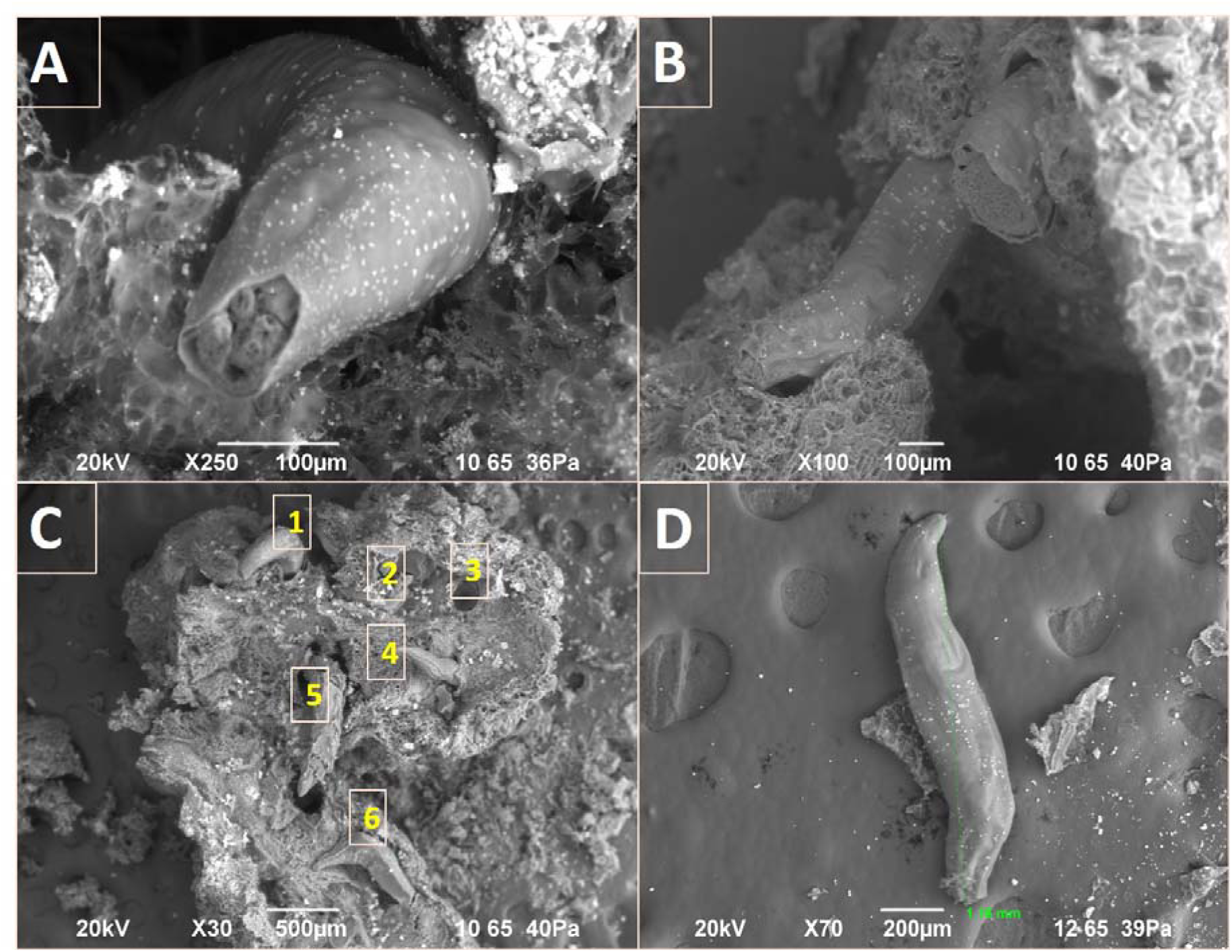
Root-knot nematodes, a) juvenile state of the nematode within the gill, wave motion within the micro-space, b) wave-like locomotion of the nematode within the micro-space of the gill, c) population of juvenile nematodes within of the knot (micro-space), d) length of a male nematode.

## Perfil elemental del knot

### Scanning electron microscopy/energy dispersive X-ray spectrometry (SEM-EDS)

We also show the elementary profile of knot infested with nematodes (Fig. 3). The elements analyzed in this study were C, N, and O. The highest to lowest concentration was C> O> N (54.73, 37.57, and 3.55%) respectively. Phosphorus (0.133%). The main cations K +, Ca2 +, Mg +, and Na2 + (0.048, 1.032, 0.094, and 1.13%). And a secondary macronutrient S (0.247%). The micronutrients and some beneficial elements Fe, Cl, Al, and Si (0.163, 0.027, 0.354, 0.985%), respectively, were also analyzed. The data, also, are corroborated by the spectra obtained (**Fig. 4**).

**Fig. 3.**
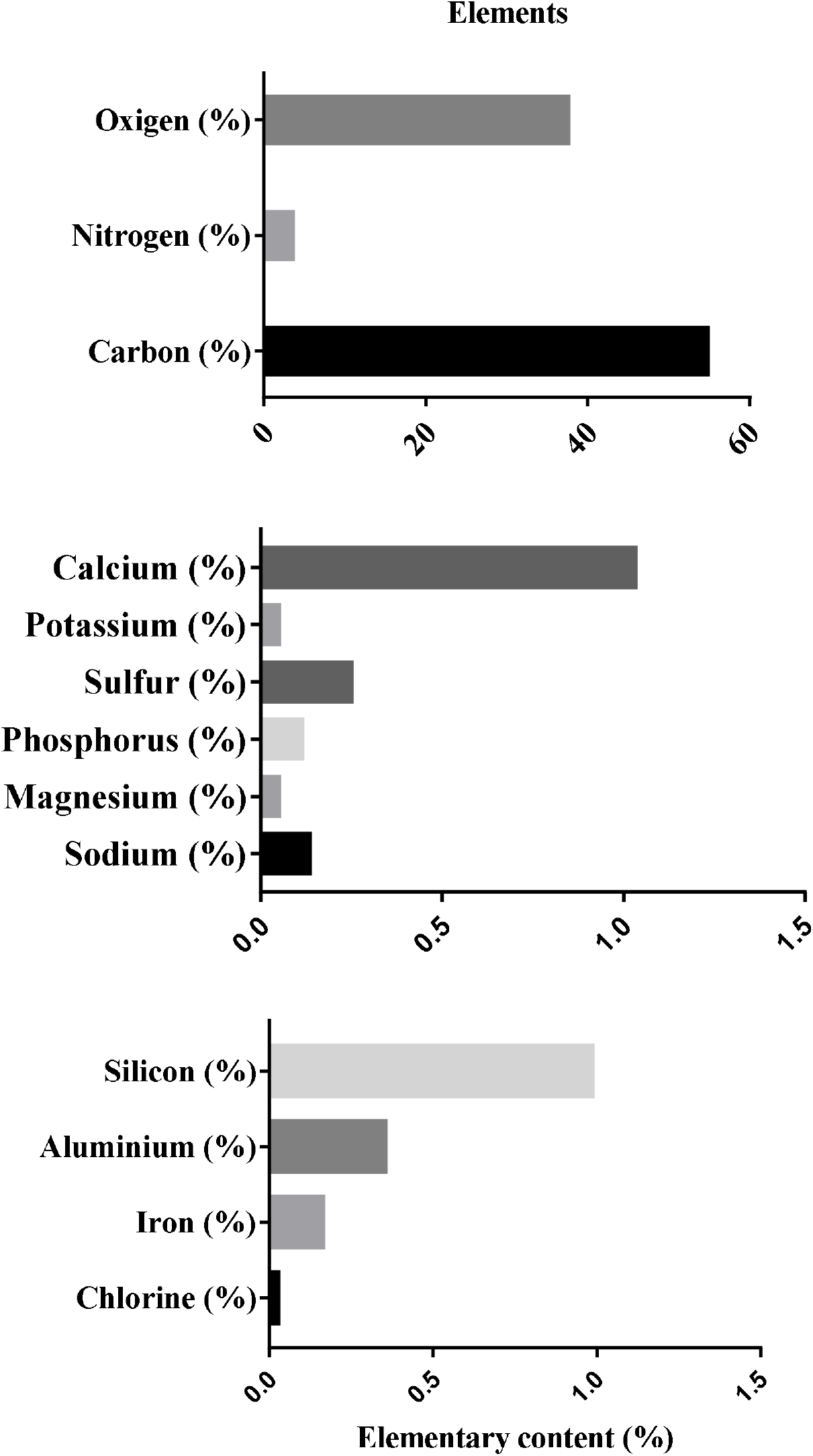
Elementary content (%) of primary and secondary macronutrients, micronutrients, and beneficial elements on root-knot nematodes.

**Fig. 4.**
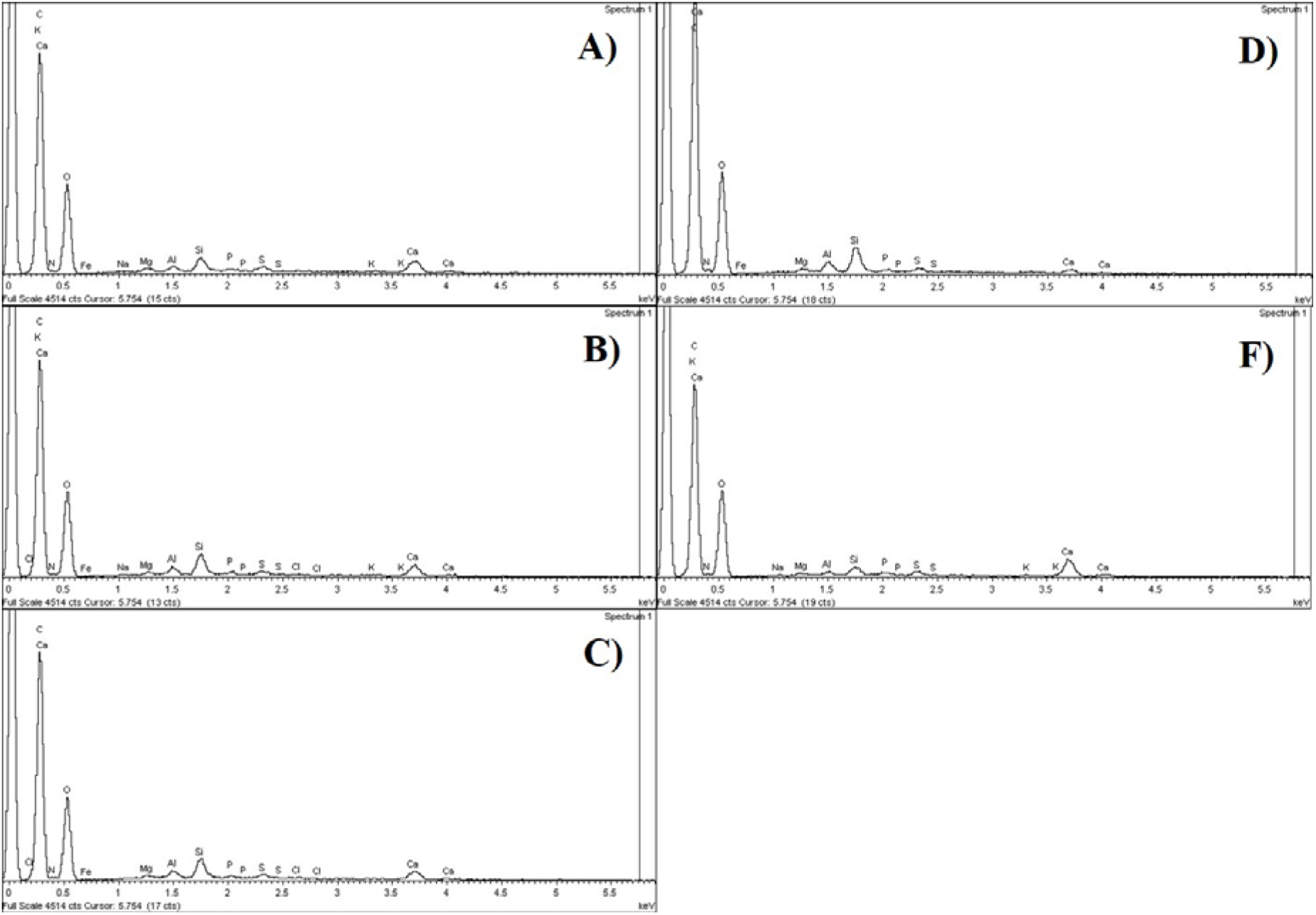
Spectra obtained from the five replicas (A, B, C, D, E, F) on (RKN). The peaks indicate the analyzed elements.

## Nematode-gall metabolic profile LC-MS Q-TOF

The chromatograms for both polarities (ES+/ES−) can be seen in Figure 5. The sample with FAA fixative solution (red peaks), presented a greater number of compounds, with respect to the sample containing HPLC water (green peak).

**Fig. 5.**
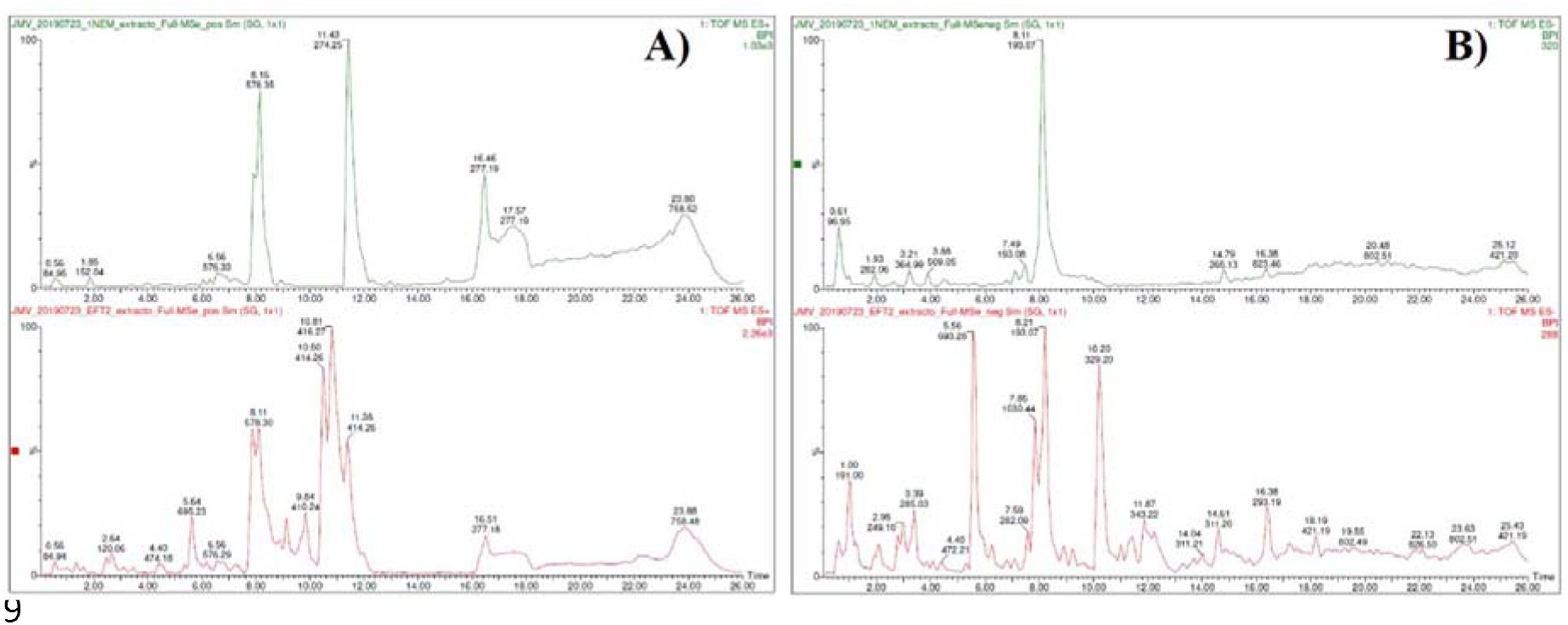
Chromatogram of progenesis. A) Polarity (ES+), the green peaks indicate the knot sample containing HPLC water; the red peaks indicate the knot sample containing FAA fixative solution. B) Polarity (ES−), the green peaks indicate the knot sample containing HPLC water; the red peaks indicate the knot sample containing FAA fixative solution.

With mode polarity (ES+) 202 compounds were detected. Of the above 195 compounds have at least one possible identification candidate, according to the database (PlantCys: https://www.plantcyc.org/) and (ChEBI: https://www.ebi.ac.uk/chebi/). While the mode (ES−) 42 compounds were detected, where 41 of the detected compounds match at least one compound. In our investigation we present the putative compounds with the highest score (Fig. 6), these were: -Oxo-1-(3-pyridyl)-1-butanone, N-Acetyl-L-tyrosine, 4-(3,4-dihydroxyphenyl) butan-2-one, Furmecyclox, Kukoamine B, Subaphylline, 2-[[3-[[2- (dimethylamino) phenyl]methyl]-2-pyridin-4-yl-1,3-diazinan-1-yl]methyl]-N,N-dimethylaniline, N1-trans-Feruloylagmatine, Physagulin E, Megestrol, Kukoamine C, Phytolaccoside A, L-3-nitrotyrosine, Egonol oleate, Collettiside I, Midodrine, Discodermolide, Dehydrotomatine, C1-[[(3S,9R,10S)-16-[[anilino(oxo)methyl]amino]-12-[(2S)-1-hydroxypropan-2-yl]-3,10-dimethyl-13-oxo-2,8-dioxa-12 azabicyclo[12.4.0]octadeca-1(14),15,17-trien-9-yl]methyl]-3-cyclohexyl-1-methylure, Dehydrotomatine, Dehydrotomatine, Jurubine, Jurubine, 17beta-Methylestra-1,3,5(10)-trien-3-ol, Veraguamide G, Etnangien, 1-palmitoyl-2-hexanoyl-sn-glycero-3- phosphocholine, Etnangien, Dehydrotomatine, Avenacin B-2, Dihydromelilotoside, (23R)-Acetoxytomatine, Tomatine, Melilotussaponin O1, Minutissamide D, (23R)-Acetoxytomatine, 4-amino-3-[(cyclohexylamino)-oxomethyl]-5 isothiazolecarboxylic acid ethyl ester, Butyl oleate sulfate, 13-hydroperoxyoctadeca-9,11-dienoic acid, 2,3-bis-O- (geranylgeranyl)-sn-glycero-3-phospho-L-serine(1-), 3-[(2S,3R)-5-[(2S)-1-hydroxypropan-2-yl]-3-methyl-2-(methylaminomethyl)-6-oxo-3,4-dihydro-2H-pyrido[2,3-b][1,5]oxazocin- 8-yl]-N,N-dimethylbenzamide. Some compounds were found more than twice in the same and different ranges (colored circles) (m/z) (Fig. 6). However, those compounds of greater agronomic interest were selected and for future research (Table 1), such as feruloylagmatine, dehydrotomatine, jurubine (Fig. 7), etnangien, tomatina, and minutissamides.

**Fig. 6.**
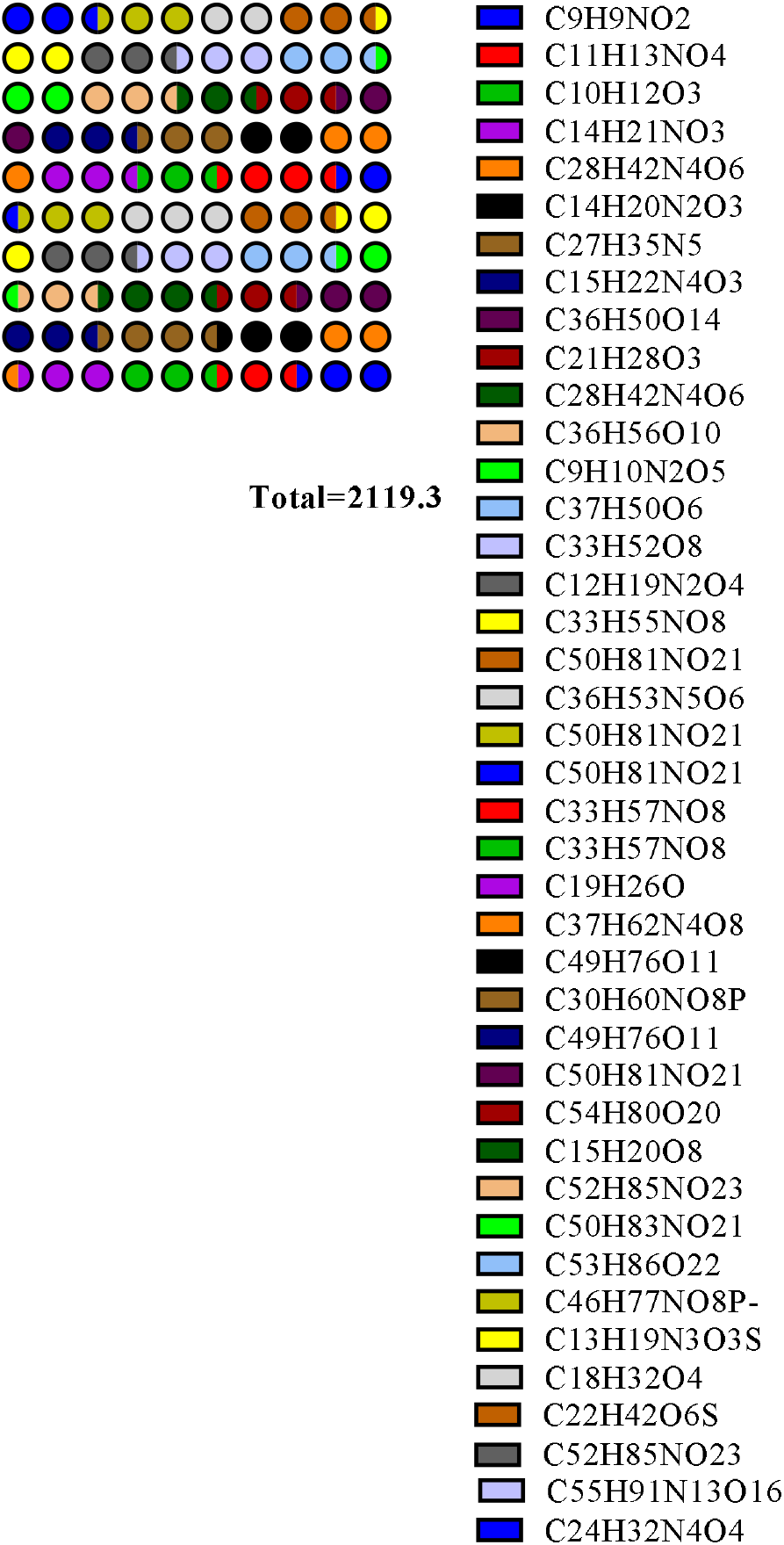
Putative metabolic compounds with the highest score found on root-knot nematodes.

**Table 1.** Compounds of interest found on (RKN) and their possible role on the root-plant system

| Molecular Formula | Fragmentation score | Mass Error (ppm) | Role of the metabolite in the root-plant | m/z |
| --- | --- | --- | --- | --- |
| C<sub>28</sub>H<sub>42</sub>N<sub>4</sub>O<sub>6</sub> | 96.1 | 1.530 | It inhibits the transduction of proinflammatory signals and the expression of cytokinins induced by LPS and CpG DNA. These are two molecular patterns associated with pathogens (PAMP). | 531.31 |
| C<sub>14</sub>H<sub>20</sub>N<sub>2</sub>O<sub>3</sub> | 95 | 2.479 | Effector of the root system of plants, influences the development of the root system and nutrient collection | 529.30 |
| C<sub>15</sub>H<sub>22</sub>N<sub>4</sub>O<sub>3</sub> | 88.3 | 1.364 | Vegetable-alkaloid metabolite, role as antifungal agent, topoisomerase DNA inhibitor. It is found in cereals and cereal products. | 307.17 |
| C<sub>14</sub>H<sub>21</sub>NO<sub>3</sub> | 95 | 2.479 | Fungicide (furamide) microbial agent that destroys fungi by suppressing its ability to grow or reproduce | 293.18 |
| C<sub>28</sub>H<sub>42</sub>N<sub>4</sub>O<sub>6</sub> | 97.2 | 1.691 | Organic compounds (catechols) | 531.31 |
| C<sub>50</sub>H<sub>81</sub>NO<sub>21</sub> | 90.8 | 1.380 | *Lycopersicon esculentum* alkaloid (steroid saponins) | 1032.39 |
| C<sub>50</sub>H<sub>81</sub>NO<sub>21</sub> | 79.9 | 0.573 | *Lycopersicon esculentum* alkaloid (steroid saponins) | 1032.53 |
| C<sub>50</sub>H<sub>81</sub>NO<sub>21</sub> | 72.2 | 1.188 | *Lycopersicon esculentum* alkaloid (steroid saponins) | 527.76 |
| C<sub>33</sub>H<sub>57</sub>NO<sub>8</sub> | 87.8 | 4.682 | It is an alkaloid that is mainly found in the fruits of *Solanum torvum* (Steroid Saponins) | 578.40 |
| C<sub>33</sub>H<sub>57</sub>NO<sub>8</sub> | 88.1 | 9.212 | It is an alkaloid that is mainly found in the fruits of *Solanum torvum* (Steroid Saponins) | 618.40 |
| C<sub>49</sub>H<sub>76</sub>O<sub>11</sub> |  |  | Macrolide antibiotic |  |
| C<sub>50</sub>H<sub>81</sub>NO<sub>21</sub> | 80.1 | 3.591 | *Lycopersicon esculentum* alkaloid (steroid saponins) | 1030.51 |
| C<sub>15</sub>H<sub>20</sub>O<sub>8</sub> | 24.1 | 3.116 | Organic compounds (phenolic glycosides). Phenolic structures include lignans and flavonoids. | 655.22 |
| C<sub>52</sub>H<sub>85</sub>NO<sub>23</sub> | 78.9 | 1.712 | *Lycoperside C* is a *Lycopersicon esculentum* alkaloid. They are organic compounds known as steroid saponins. | 1090.54 |
| C<sub>50</sub>H<sub>83</sub>NO<sub>21</sub> | 90.8 | 3.142 | Glucoalkaloid (tomatidine) has a role as a fungal and antimicrobial immune adjuvant. It is found in tomato leaves. | 1078.54 |
| C<sub>55</sub>H<sub>91</sub>N<sub>13</sub>O<sub>16</sub> | 93.5 | 0.446 | Organic compounds triterpenic saponins. They are glycosylated derivatives of triterpenic sapogenins | 1095.53 |

**Fig. 7.**
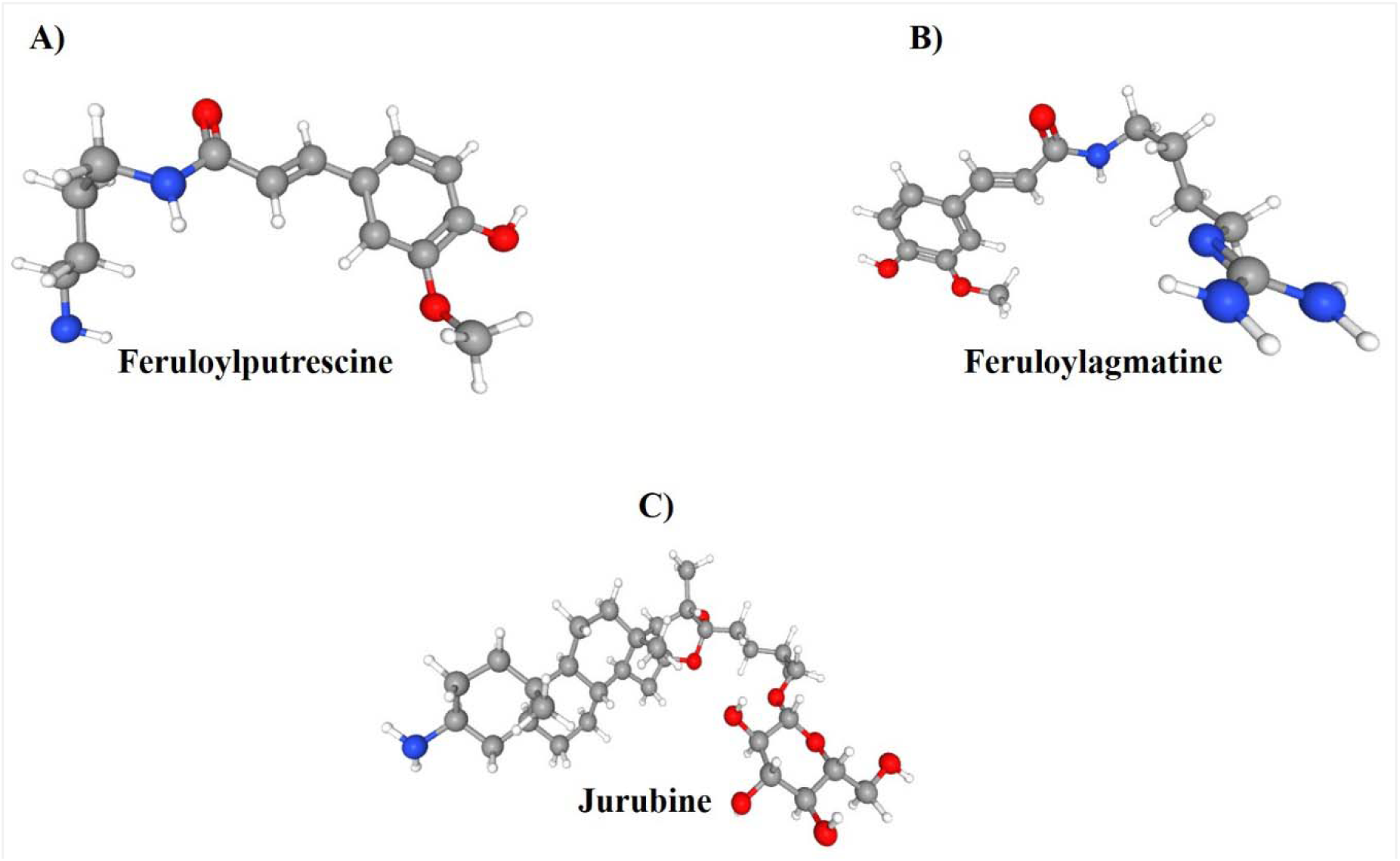
Compounds of great agronomic interest on root-knot nematodes. A) Feruloylputrescina soluble conjugated polyamide (C_14_H_20_N_2_O_3_) is related to the colonization of arbuscular mycorrhizal fungi on the knot, it can be induced by intraradic *Glomus* and it can have a positive response to biotic and abiotic stress. B) Feruloylagmatine C_15_H_22_N_4_O_3_ is an alkaloid that inhibits the germination of spores of *Fusarium* pseudograminearum. B) The jurubine (C33H57NO8) is an alkaloid with antioxidant properties, and has been reported in *Solanum torvum* and is synthesized in leaves, roots and fruits.

## Discussion

### Pathogenesis: Root-knot nematodes (SEM-EDS)

Nematodes in juvenile state were observed for the first time in the micro-spaces within the gill. The wave motion is corroborated with the micrographs (Fig. 2B), described studies of wave movements in Newtonian fluids were described for *C. elegans* (Fang-Yen et al., 2010).

The individual giant cells induced by the nematodes, is the perfect microspace for their proliferation and feeding (Fig. 2C). After embryogenesis, the juvenile of the first stage (J1) moves into the egg, and passes to the infectious juvenile of the second stage (J2) that emerges from the egg (Jones *et al*., 2013). The damage is due to the diversion of the host’s nutrients to the knot and interference with transport processes (Chin *et al*., 2018). Plants can recognize cell damage caused by invasion of parasitic nematodes (PPN), through recognition based on patterns located on the cell surface (PRR) of molecular patterns associated with damage (Sato *et al*., 2019).

### Elementary relative concentration on root-knot nematodes (SEM-EDS)

The roots absorb water, mineral nutrients and anchor to the plant; outbreaks perform photosynthesis, perspiration, and are the breeding site (Groff and Kaplan, 1988). The relative elemental concentration on the gill was C>O>N (54.73, 37.57, and 3.55%) respectively. Phosphorus (0.133%). The main cations Na>Ca>Mg>K (1.13, 1.032, 0.094 and 0.048%). The S as secondary macronutrient (0.247%). The micronutrients were Fe>Cl (0.163, 0.027%), and the beneficial elements Si>Al (0.985, 0.354%) respectively. The data, also, are corroborated by graph and spectra obtained (Fig. 3, Fig. 4). C, H, and O, are the non-mineral class of macronutrients, however, plants depend on them for the construction of larger organic molecules in the cell. Macronutrients are related to cellular components, such as proteins and nucleic acids, are required in large quantities. Micronutrients are the important enzyme cofactors in plant metabolism and are required in small amounts (Morgan and Connolly, 2013). Roots >2 mm in diameter that contained significantly more N, P, and Mg and less C were analyzed, the concentrations for N and P were 11.0 and 0.9 g·kg respectively, more than the 5 mm roots of coniferous trees and broad leaf (Gordon and Jackson, 2000). This indicates that the distribution of minerals in the different organs of the plant may vary over time (Juárez-Maldonado *et al*., 2017).

### Metabolic signal transduction on (RKN)

Feeding sites serve as nutrient stores for nematodes, but it is in stark contrast, with the mutual benefits derived from microbes, such as plants during mycorrhizal root colonization (Davis and Mitchum, 2005). Genes encoding secreted chorismate mutase (CM) have been isolated from root-knot and soybean cyst nematodes (Doyle and Lambert, 2003). CM is a pivotal enzyme in the shikimic acid pathway that modulates synthesis of Phe and Try, having pleiotropic effects on cellular metabolism, auxin synthesis, and as precursors of plant defense compounds (Davis and Mitchum, 2005). In this first metabolomic profile, kukoamine (KB, C_28_H_42_N_4_O_6_) (Table 1) was detected, a natural alkaloid with high affinity for bacterial lipopolysaccharide DNA (PLS) and oligodeoxynucleotides containing CpG motifs (Liu *et al*., 2011). The metabolic profile detected fungicidal molecules such as furmecyclox C_14_H_21_NO_3_ possibly for root rot control (Kataria and Verma, 1993). The soluble conjugated polyamide feruloylputrescin (C_14_H_20_N_2_O_3_) is reported for the first time (Tang and Newton, 2005) (Table 1, Fig. 7). The above polyamide may be related to colonization of arbuscular mycorrhizal fungi on (RKN). This accumulation of metabolites can be induced by fungi such as *Glomus intraradices* (Peipp *et al*., 1997). In addition, this polyamide has a critical role in response to abiotic stress and defense system against pathogens (Pihlava, 2014).

Feruloylagmatine C_15_H_22_N_4_O_3_ is an alkaloid (Table 1, Fig. 7) of the guanidine type belonging to the Papaveraceae family (Cheng *et al*., 2008). This compound seems to inhibit the germination of *Fusarium pseudograminearum* spores, but they have no effect on mycelial growth in the *Brachypodium distachyon* Bdact2a mutant (Carere *et al*., 2018). Its biosynthesis is higher in roots than in barley sprouts, this pattern could be used to regulate antifungal properties (Gorzolka *et al*., 2014). Tomato plants synthesize glycoalcaloid dehydrotomatine (C_50_H_81_NO_21_) and alpha-tomatine, these glycoalcaloids possibly act as a defense with bacteria, fungi, viruses and insects, and have been found at different stages of maturity, calyces, flowers, leaves, roots and stems (Kozukue *et al*., 2004). Toxic compounds of dehydrotomatine in foliage and immature fruits prevent diseases in plants (Cataldi *et al*., 2005). Potatoes, too, produce these glycoalkaloids and biologically active metabolites that have specific roles in plants and the human diet (Friedman, 2006). This bioactivity of glycoalkaloids in the genus *Solanum*, include its anticancer, anticolesterol, antimicrobial, anti-inflammatory, antinociceptive, and antipyretic effect (Milner *et al*., 2011). We report for the first time the jurubine compound (C_33_H_57_NO_8_) (Table 1, Fig. 7) in the metabolic profile of the nematode knot, an alkaloid with antioxidant properties, and has been reported in *Solanum torvum* (Innih *et al*., 2018). It is known that the genus *Solanum* synthesize this compound in leaves, roots and fruits (Scheiber and Ripperger, 1996). We report for the first time a novel natural macrolide antibiotic called etnangien (C_49_H_76_O_11_) (Table 1), the fact is that it is a potent structurally unique RNA polymerase inhibitor analogue of myxobacteria (*Soragium cellulosum*) (Menche *et al*., 2008). This gram-negative bacterium secretes hydrolytic enzymes that break down organic matter (OM) and other living organisms in its environment, in addition, they form latent fruiting bodies that resist environmental stresses (Julien and Fehd, 2003). These findings demonstrate the relationship between the nematode-knot-nutrients- and possible myxobacteria in tomato roots.

We report, to the compound dihydromelilotosida C_15_H_20_O_8_ (Table 1), this metabolite has been observed in Dendrobium, a compound with antioxidant potential (Yang et al., 2007; Gutiérrez, 2010), 56 compounds have been reported in the flower structure of the *Cymbidium* orchid with different functionalities (García-Gaytán et al., 2013). Tomatine (C_50_H_83_NO_21_), like dehydrotomatin, is stereroid glycoalkaloids (SGA), and is located in immature leaves and fruits of tomatoes, it is possible that this alkaloid protects the fruits while they ripen (Caballero *et al*., 2003; Cárdenas *et al*., 2016). In the metabolomic profile we find the compound minutissamides (C_55_H_91_N_13_O_16_) (Table 1), a cyclic polypeptide that possess cytotoxic activity (Mareš *et al*., 2019). Four A-D minutissamide of the *Anabaena minutissma* cyanobacterium (UTEX 1613) have been isolated (Kang *et al*., 2011). Facing the challenges of water resources in the world, he has investigated cyanobacteria, they have the biological potential for wastewater treatment, there is a 70% reduction in calcium, 46% chloride, 100% nitrate, 88% nitrite, 100% ammonia, 92 % total phosphorus, 12.5% magnesium in different wastewater treated with different species of cyanobacteria (Dash *et al*., 2019). Cyanobacteria have the ability to switch to myxotrophic nutrition, they can metabolize organic pollutants, breaking them down into less toxic or non-toxic substances (bio-remediation), while in the purification of wastewater it is based on their ability to use ammonium ions as nitrogen source (Zinicovscaia and Cepoi, 2016).

## Conclusion

Root-knot nematodes are parasites that destroy many crops globally. We adopt for the first time, the term of micro-space of the nematode within the knot. In all cases the nematodes were found in infectious juvenile state (J2), with wave motion. The presence of Na^+^ on root-knot nematodes affected the concentration of Mg^2+^ and K^+^, while the relative concentration of N was higher. Possibly this response is due to the type of fertilizer used and/or nutrient solution. Tuna et al. (2007) found that the Ca/Na and K/Na ratio can be affected in tomato leaves, due to a higher concentration of NaCl in the root, and the concentration of N is not affected. The most relevant compounds for future research are kokoamine, furmecyclox, feruloylputrescina, feruloylagmatina, dehydrotomatina, jurubine, etnangien, dehydromelilotoside, tomatina and minutissamides. It is suggested that an integrated management and control over root-knot nematodes, should be carried out, laboratory analysis of soil, and water. The application of composts and vermicomposts on planting beds, mineral fertilizers, control of the nutrient solution (EC, pH), foliar applications of biostimulants, monitoring of relative humidity (RH) and temperature for pest control/diseases.

## Supplementary data

**Supplementary data are available at JXB online**

**Figure S1.**Tomato root nematodes seen from the stereoscope microscope

## Acknowledgments

To Dr. Emanuel Bojórquez-Quintal (Professor-CONACTY) for the conservation and treatment of the sample. Mr. Esteban Sánchez-Rodríguez (Colmich-LADIPA) for taking the micrographs and determining the elementary content on (RKN). Dr. José J. Ordaz-Ortiz and Mr. Josaphat M. Montero Vargas (Lab metabolomics: Metabolomics and mass spectrometry, Cinvestav-Langebio) for the methodology, elaboration of the analysis and results of the metabolomic profile on (RKN). Olimpia Alonso-Pérez analyst (Colmich-LADIPA). Analyst Luis M. Rojas-Abarca (Colmich-LADIPA). Students Elena Gómez-Cabrera and Juan L. Negrete-Guerrero (UPP) for their support, and Dr. Víctor García-Gaytán idea, writing, statistical analysis, interpretation, revision, and corrections of the manuscript (Research Professor, El Colegio de Michoacán, AC).

